# Genetics of brain age suggest an overlap with common brain disorders

**DOI:** 10.1101/303164

**Authors:** Tobias Kaufmann, Dennis van der Meer, Nhat Trung Doan, Emanuel Schwarz, Martina J. Lund, Ingrid Agartz, Dag Alnæs, Deanna M. Barch, Ramona Baur-Streubel, Alessandro Bertolino, Francesco Bettella, Mona K. Beyer, Erlend Bøen, Stefan Borgwardt, Christine L. Brandt, Jan Buitelaar, Elisabeth G. Celius, Simon Cervenka, Annette Conzelmann, Aldo Córdova-Palomera, Anders M. Dale, Dominique J.-F de Quervain, Pasquale Di Carlo, Srdjan Djurovic, Erlend S. Dørum, Sarah Eisenacher, Torbjørn Elvsåshagen, Thomas Espeseth, Helena Fatouros-Bergman, Lena Flyckt, Barbara Franke, Oleksandr Frei, Beathe Haatveit, Asta K. Håberg, Hanne F. Harbo, Catharina A. Hartman, Dirk Heslenfeld, Pieter J. Hoekstra, Einar A. Høgestøl, Terry Jernigan, Rune Jonassen, Erik G. Jönsson, Karolinska Schizophrenia Project (KaSP), Peter Kirsch, Iwona Kłoszewska, Knut-Kristian Kolskår, Nils Inge Landrø, Stephanie Le Hellard, Klaus-Peter Lesch, Simon Lovestone, Arvid Lundervold, Astri J. Lundervold, Luigi A. Maglanoc, Ulrik F. Malt, Patrizia Mecocci, Ingrid Melle, Andreas Meyer-Lindenberg, Torgeir Moberget, Linn B. Norbom, Jan Egil Nordvik, Lars Nyberg, Jaap Oosterlaan, Marco Papalino, Andreas Papassotiropoulos, Paul Pauli, Giulio Pergola, Karin Persson, Geneviève Richard, Jaroslav Rokicki, Anne-Marthe Sanders, Geir Selbæk, Alexey A. Shadrin, Olav B. Smeland, Hilkka Soininen, Piotr Sowa, Vidar M. Steen, Magda Tsolaki, Kristine M. Ulrichsen, Bruno Vellas, Lei Wang, Eric Westman, Georg C. Ziegler, Mathias Zink, Ole A. Andreassen, Lars T. Westlye, for the Alzheimer’s Disease Neuroimaging Initiative, for the Pediatric Imaging, Neurocognition and Genetics Study, for the AddNeuroMed consortium

**Affiliations:** NORMENT, KG Jebsen Centre for Psychosis Research, Division of Mental Health and Addiction, Oslo University Hospital & Institute of Clinical Medicine, University of Oslo, Oslo, Norway; Department of Psychiatry and Psychotherapy, Central Institute of Mental Health, Medical Faculty Mannheim, Heidelberg University, Mannheim, Germany; Department of Psychiatry, Diakonhjemmet Hospital, Oslo, Norway; Department of Clinical Neuroscience, Center for Psychiatry Research, Karolinska Institutet and Stockholm County Council, Stockholm, Sweden; Department of Psychological and Brain Sciences, Washington University in St. Louis, St. Louis, USA; Department of Psychiatry, Washington University in St. Louis, St. Louis, USA; Department of Radiology, Washington University in St. Louis, St. Louis, USA; Department of Psychology I, University of Würzburg, Würzburg, Germany; Institute of Psychiatry, Bari University Hospital, Bari, Italy; Department of Basic Medical Science, Neuroscience, and Sense Organs, University of Bari, Bari, Italy.; Institute of Clinical Medicine, University of Oslo, Oslo, Norway; Department of Radiology and Nuclear Medicine, Section of Neuroradiology, Oslo University Hospital, Oslo, Norway; Department of Psychiatry (UPK), University of Basel, Basel, Switzerland; Institute of Psychiatry, King’s College, London, UK; Department of Cognitive Neuroscience, Donders Institute for Brain, Cognition and Behaviour, Radboud University Medical Center, Nijmegen, The Netherlands; Karakter Child and Adolescent Psychiatry University Centre, Nijmegen, The Netherlands; Institute of Health and Society, University of Oslo, Oslo, Norway; Department of Neurology, Oslo University Hospital, Oslo, Norway; Children and Adolescence Psychiatry, University of Tübingen, Tübingen, Germany; Department of Pediatrics, Stanford University School of Medicine, Stanford University, Stanford, USA; Department of Radiology, University of California, San Diego, La Jolla, CA, USA; Department of Neurosciences, University of California, San Diego, La Jolla, CA, USA; Division of Cognitive Neuroscience, University of Basel, Basel, Switzerland; Transfaculty Research Platform Molecular and Cognitive Neurosciences, University of Basel, Basel, Switzerland; Department of Medical Genetics, Oslo University Hospital, Oslo, Norway; NORMENT, K.G. Jebsen Center for Psychosis Research, Department of Clinical Science, University of Bergen, Bergen, Norway; Department of Psychology, University of Oslo, Oslo, Norway; Sunnaas Rehabilitation Hospital HT, Nesodden, Norway; Departments of Human Genetics and Psychiatry, Donders Institute for Brain, Cognition and Behaviour, Radboud University Medical Center, Nijmegen, The Netherlands; Department of Neuromedicine and Movement Science, Norwegian University of Science and Technology, Trondheim, Norway; Department of Radiology and Nuclear Medicine, St. Olavs Hospital, Trondheim, Norway; Department of Psychiatry, University of Groningen, University Medical Center Groningen, Groningen, The Netherlands; Department of Clinical Neuropsychology, Vrije Universiteit Amsterdam, Amsterdam, The Netherlands; Department of Cognitive Psychology, Vrije Universiteit Amsterdam, Amsterdam, The Netherlands; Department of Child and Adolescent Psychiatry, University Medical Center Groningen, University of Groningen, Groningen, The Netherlands; Center for Human Development, University of California, San Diego, USA; Department of Cognitive Science, University of California, San Diego, USA; Departments of Psychiatry and Radiology, University of California, San Diego, USA; Members of Karolinska Schizophrenia Project (KaSP) are listed before References; Department of Clinical Psychology, Central Institute of Mental Health, Medical Faculty Mannheim, Heidelberg University, Mannheim, Germany; Bernstein Center for Computational Neuroscience Heidelberg/Mannheim, Mannheim, Germany; Department of Old Age Psychiatry and Psychotic Disorders, Medical University of Lodz, Poland; Division of Molecular Psychiatry, Center of Mental Health, University of Würzburg, Würzburg, Germany; Laboratory of Psychiatric Neurobiology, Institute of Molecular Medicine, Sechenov First Moscow State Medical University, Moscow, Russia; Department of Neuroscience, School for Mental Health and Neuroscience (MHeNS), Maastricht University, Maastricht, The Netherlands; Department of Psychiatry, Warneford Hospital, University of Oxford, Oxford, UK; Department of Biomedicine, University of Bergen, Norway; Mohn Medical Imaging and Visualization Centre, Department of Radiology, Haukeland University Hospital, Bergen, Norway; Department of Biological and Medical Psychology, University of Bergen, Norway; K. G. Jebsen Centre for Neuropsychiatric Disorders, University of Bergen, Norway; Department of Research and Education, Oslo University Hospital, Oslo, Norway; Institute of Gerontology and Geriatrics, University of Perugia, Perugia, Italy; Department of Radiation Science, Umeå University, Umeå, Sweden; Emma Children’s Hospital Amsterdam Medical Center, Amsterdam, The Netherlands; VU University Medical Center, Department of Pediatrics, Amsterdam, The Netherlands; Division of Molecular Neuroscience, University of Basel, Basel, Switzerland; Life Sciences Training Facility, Department Biozentrum, University of Basel, Basel, Switzerland; Department of Geriatric Medicine, Oslo University Hospital, Oslo, Norway; Norwegian National Advisory Unit on Ageing and Health, Vestfold Hospital Trust, Tønsberg, Norway; Centre for Old Age Psychiatric Research, Innlandet Hospital Trust, Ottestad, Norway; Department of Neurology, Institute of Clinical Medicine, University of Eastern Finland, Kuopio, Finland; Neurocenter, Neurology, Kuopio University Hospital, Kuopio, Finland; Dr. E. Martens Research Group for Biological Psychiatry, Department of Medical Genetics, Haukeland University Hospital, Bergen, Norway.; 1^st^ Department of Neurology, Aristotle University of Thessaloniki, Makedonia, Greece.; INSERM U 1027, University of Toulouse, Toulouse, France; Feinberg School of Medicine, Northwestern University, Chicago, Illinois, USA; Department of Neurobiology, Care Sciences and Society, Karolinska Institute, Stockholm, Sweden; District hospital Ansbach, Germany; Data used in preparation of this article were obtained from the Alzheimer’s Disease Neuroimaging Initiative (ADNI) database (adni.loni.usc.edu). As such, the investigators within the ADNI contributed to the design and implementation of ADNI and/or provided data but did not participate in analysis or writing of this report. A complete listing of ADNI investigators can be found at: http://adni.loni.usc.edu/wp-content/uploads/how_to_apply/ADNI_Acknowledgement_List.pdf.; Data used in preparation of this article were obtained from the Pediatric Imaging, Neurocognition and Genetics Study (PING) database (http://ping.chd.ucsd.edu). As such, the investigators within PING contributed to the design and implementation of PING and/or provided data but did not participate in analysis or writing of this report. A complete listing of PING investigators can be found at https://ping-dataportal.ucsd.edu/sharing/Authors10222012.pdf; AddNeuroMed consortium was led by Simon Lovestone, Bruno Vellas, Patrizia Mecocci, Magda Tsolaki, Iwona Kłoszewska, Hilkka Soininen

**Keywords:** Brain age gap, brain disorders, genetic architecture, pleiotropy

## Abstract

Numerous genetic and environmental factors contribute to psychiatric disorders and other brain disorders. Common risk factors likely converge on biological pathways regulating the optimization of brain structure and function across the lifespan. Here, using structural magnetic resonance imaging and machine learning, we estimated the gap between brain age and chronological age in 36,891 individuals aged 3 to 96 years, including individuals with different brain disorders. We show that several disorders are associated with accentuated brain aging, with strongest effects in schizophrenia, multiple sclerosis and dementia, and document differential regional patterns of brain age gaps between disorders. In 16,269 healthy adult individuals, we show that brain age gap is heritable with a polygenic architecture overlapping those observed in common brain disorders. Our results identify brain age gap as a genetically modulated trait that offers a window into shared and distinct mechanisms in different brain disorders.

---

Psychiatric disorders and other brain disorders are among the main contributors to morbidity and disability around the world^1^, often placing debilitating disadvantages on the individual^2^. The disease mechanisms are complex, spanning a wide range of genetic and environmental contributing factors^3^. The inter-individual variability is large, but on a group-level, patients with common brain disorders perform worse on cognitive tests, are less likely to excel professionally, and engage in adverse health behaviours more frequently^4^.

Dynamic processes influencing the rate of maturation and change throughout the lifespan play a critical role, as reflected in the wide range of disease onset times from early childhood to old age^5^. This suggests that the age at which individual trajectories diverge from the norm is a key characteristic in the underlying pathophysiology. Whereas autism spectrum disorder (ASD) and attention-deficit/hyperactivity disorder (ADHD) have onset in childhood^6^, schizophrenia (SZ) and bipolar (BD) spectrum disorders likely evolve during adolescence, before the outbreak of severe symptoms, which is typically in early adulthood^6,7^. Likewise, multiple sclerosis (MS) most often presents itself in early adulthood but the disease process likely starts much earlier^8,9^. First episodes in major depressive disorder (MDD) can appear at any stage from adolescence to old age^6,10^, whereas mild cognitive impairment (MCI) and dementia (DEM) most often evolve in old age^11^. When attempting to decode the underlying brain dysfunction of these disorders, age-related deviations from the norm may also differ in terms of spatial location, direction, change rate and magnitude, all of which add complexity to the interpretation of observed effects.

Magnetic resonance imaging (MRI) is a powerful tool to unveil abnormal brain development^12,13^ and age-related degeneration^14,15^. Machine learning techniques enable robust estimation of the biological age of the brain using MRI-derived features^16^, and initial evidence suggests that a deviation between brain and chronological age – termed the *brain age gap* - is present in several brain disorders^17^. These findings may render brain age gap a promising marker of brain health^17^, but several critical issues remain to be addressed. First, while advantageous for narrowing the complexity, reducing a rich set of brain imaging features into a single estimate of brain age inevitably compromises spatial specificity, thereby potentially removing disorder-specific patterns. Second, most studies so far have been rather small-scale, performed within a limited age range and focusing on a single disorder, which left them unable to uncover clinical specificity and lifespan dynamics. Third, the genetic underpinnings of brain age gap are not understood and it is unknown to what degree they overlap with the genetic architecture of major clinical traits. To address these critical knowledge gaps, large imaging genetics samples covering a range of prevalent brain disorders are necessary.

The availability of unprecedented sample sizes of neuroimaging and genetics data through global data sharing and population-based efforts provide new opportunities for accurate modelling of lifespan differences in brain anatomy and its application to brain disorders^18^. Here, we have gathered raw structural MRI data from a large number of individuals. We employed a centralized and harmonized processing protocol including automated surface-based morphometry and subcortical segmentation using Freesurfer^19^ (Suppl. Fig. 1). The main analysis in this study is based on data from 36,891 individuals aged 3 to 96 years that passed quality control, representing the largest brain imaging study on brain age to date.

This sample included data from healthy controls (HC; *n* = 30,967; 3-95 years), as well as from 5924 individuals with diverse brain disorders with typical onset age distributed across the lifespan^6,11^. We included data from individuals with ASD (*n* = 975; 5-64 years) and ADHD (*n* = 751; 7-62 years), individuals with prodromal SZ or at risk mental state (SZRISK; *n* = 98; 16-42 years), individuals with SZ (n = 1,145; 18-66 years), a heterogeneous group with mixed diagnoses in the psychosis spectrum (PSYMIX; *n* = 294; 18-69 years), individuals with BD (*n* = 445; 18-66 years), MS (*n* = 254; 19-68 years), MDD (*n* = 211; 18-71 years), MCI (*n* = 992; 38-91 years), and DEM (including Alzheimer’s disease (AD); *n* = 759; 53-96 years). Supplementary Tables 1 and 2 provide details on the sample’s characteristics.

## Brain age prediction across brain disorders

We used machine learning to estimate individual biological brain age based on structural brain imaging features. First, we grouped all subjects into different samples. For each of the ten clinical groups, we identified a group of healthy individuals of equal size, matched on age, sex and scanning site using propensity score matching^20^. All remaining individuals were joined into one sample comprising healthy individuals only. The latter constituted a training sample, used to train and tune the machine learning models for age prediction (*n* = 26,535; 14,182 females and 12,353 males, aged 3-89 years), whereas the ten clinical samples were used as independent test samples. Figure 1a illustrates the respective age distributions per sex and diagnosis.

**Figure 1:**
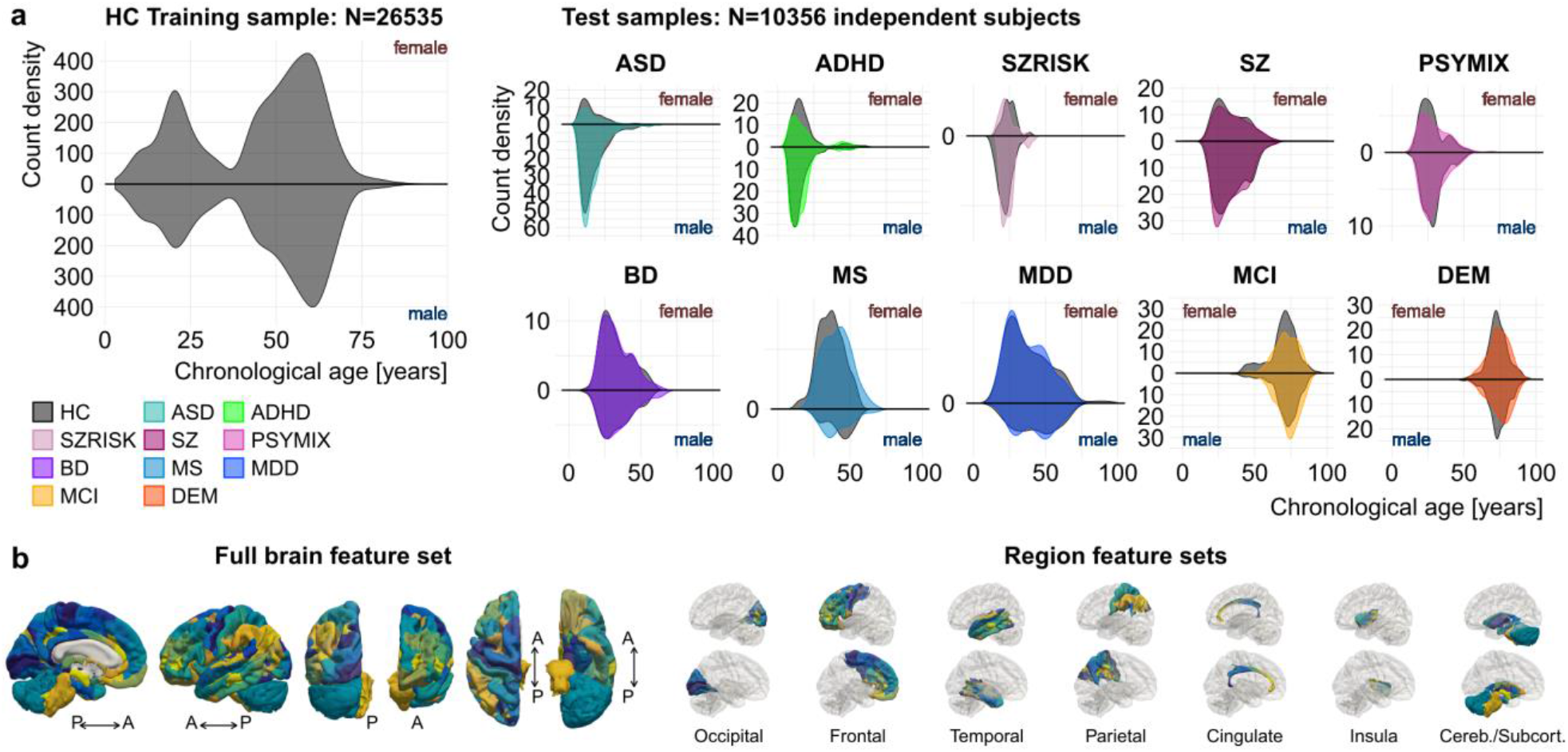
Sample distributions and brain features used for brain age prediction. **a**, Age distributions of the training (left) and the ten test samples (right) per sex and diagnosis. The groups in each test sample were of equal size and were matched for age, sex and scanning site^20^. **b**, Cortical features from the Human Connectome Project (HCP) atlas^21^ as well as cerebellar/subcortical features^19^ used for brain age prediction. Colours were assigned randomly to each feature. All features were used in the full brain feature set (left), whereas only those from specific regions (occipital, frontal, temporal, parietal, cingulate, insula, cerebellar/subcortical) were included in the region-wise feature set (right). For illustration purpose, the left hemisphere is shown.

The large sample size and wide age-span of the training sample allowed us to model male and female brain age separately, thereby accounting for potential sexual dimorphisms in brain structural lifespan trajectories. For each sex, we built a machine learning model based on gradient tree boosting (*xgboost*)^22^ to predict the age of the brain from a set of thickness, area and volume features extracted using a multi-modal parcellation of the cerebral cortex^21^ as well as a set of cerebellar/subcortical volume features^19^ (1,118 features in total, Fig. 1b). Five-fold cross-validations confirmed the validity of the models, yielding high correlations between chronological age and predicted brain age (r=.94 and r=.95 for the female and male model, respectively; Suppl. Fig. 2). Next, we applied the models to predict brain age for each individual in the ten independent test samples, and tested for effects of diagnosis on the brain age gap. We used mega-analysis (across-site analysis) as the main statistical framework as it may best exploit the benefits of the big data approach, while also providing results from a meta-analysis framework in the supplement. We controlled all associations and group differences reported in this paper for age, age², sex, and scanning site. Further, to rule out confounding effects of data quality on the results^23^, we repeated the main analyses using a more stringent multivariate quality control and exclusion procedure^24^.

Figure 2 illustrates that the brain age gap was increased in several brain disorders. Strongest effects were observed in SZ (Cohen’s *d* = 0.55), MS (*d* = 0.72), MCI (*d* = 0.45) and DEM (*d* = 1.06). PSYMIX (*d* = 0.22) and BD (*d* = 0.30) showed small effects of increased brain age gap, whereas other groups showed negligible effects. The meta-analysis converged on the same findings (Suppl. Fig. 3) and the results replicated regardless of the quality control exclusion criterion applied (Suppl. Fig. 4). Compared to matched healthy controls, the average brain age gap was estimated to 1.1 years for ASD, 0.7 years for ADHD, 0.6 years for SZRISK, 3.9 years for SZ, 1.4 years for PSYMIX, 2.0 years for BD, 5.6 years for MS, 0.8 years for MDD, 3.0 years for MCI and 5.8 years for DEM. The brain age gap in all clinical groups was positive and there were no signs of a negative brain age gap (delay) in children with ASD or ADHD.

**Figure 2:**
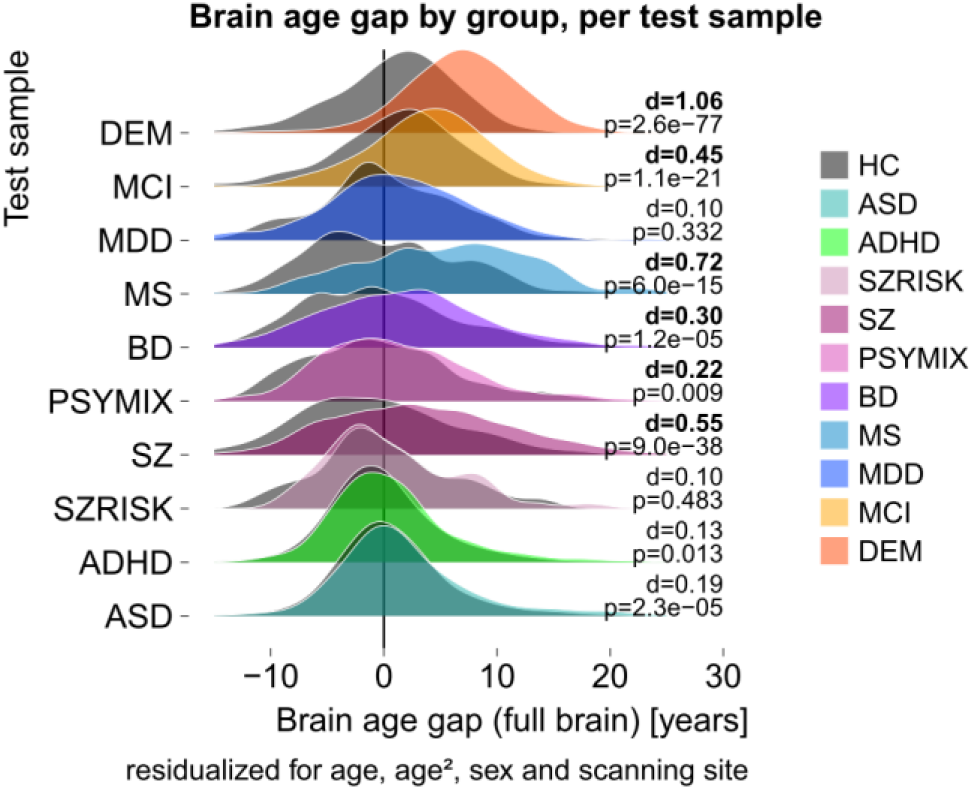
Accentuated brain aging is common in several brain disorders. Compared to healthy controls matched for age, sex and scanning site, the gap between chronological age and brain age was increased in several disorders. Cohen’s d effect sizes indicate largest effects in SZ, MS, MCI and DEM.

## Regional specificity of brain age gap

We assessed the specificity of the spatial brain age gap patterns across clinical groups. We trained age prediction models similar to those for the full brain above, including only occipital, frontal, temporal, parietal, cingulate, insula, or cerebellar/subcortical features (Fig. 1b). Cross-validation confirmed the predictive performance of all regional models (Suppl. Fig. 2), so we used these to predict regional brain age in the ten independent test sets. Region-wise brain age gaps often corresponded to the ones observed on the full brain level, yet some notable differences in the spatial patterns of the disorders emerged (Fig. 3a). For example, increased cerebellar/subcortical age gap was most prominent in DEM (*d* = 0.99) and MS (*d* = 0.89) but was not present in SZ (*d* = 0.08). The largest effect in SZ was observed in the frontal lobe (*d* = 0.71). In PSYMIX, brain age gaps in the insula (*d* = 0.39) and the temporal lobe (*d* = 0.34) were larger than the brain age gap observed on the full brain level (*d* = 0.22). A brain age gap in the temporal lobe was observed in MDD (*d* = 0.26), whereas there was no evidence for a brain age gap in ASD or ADHD in any of the regions.

**Figure 3:**
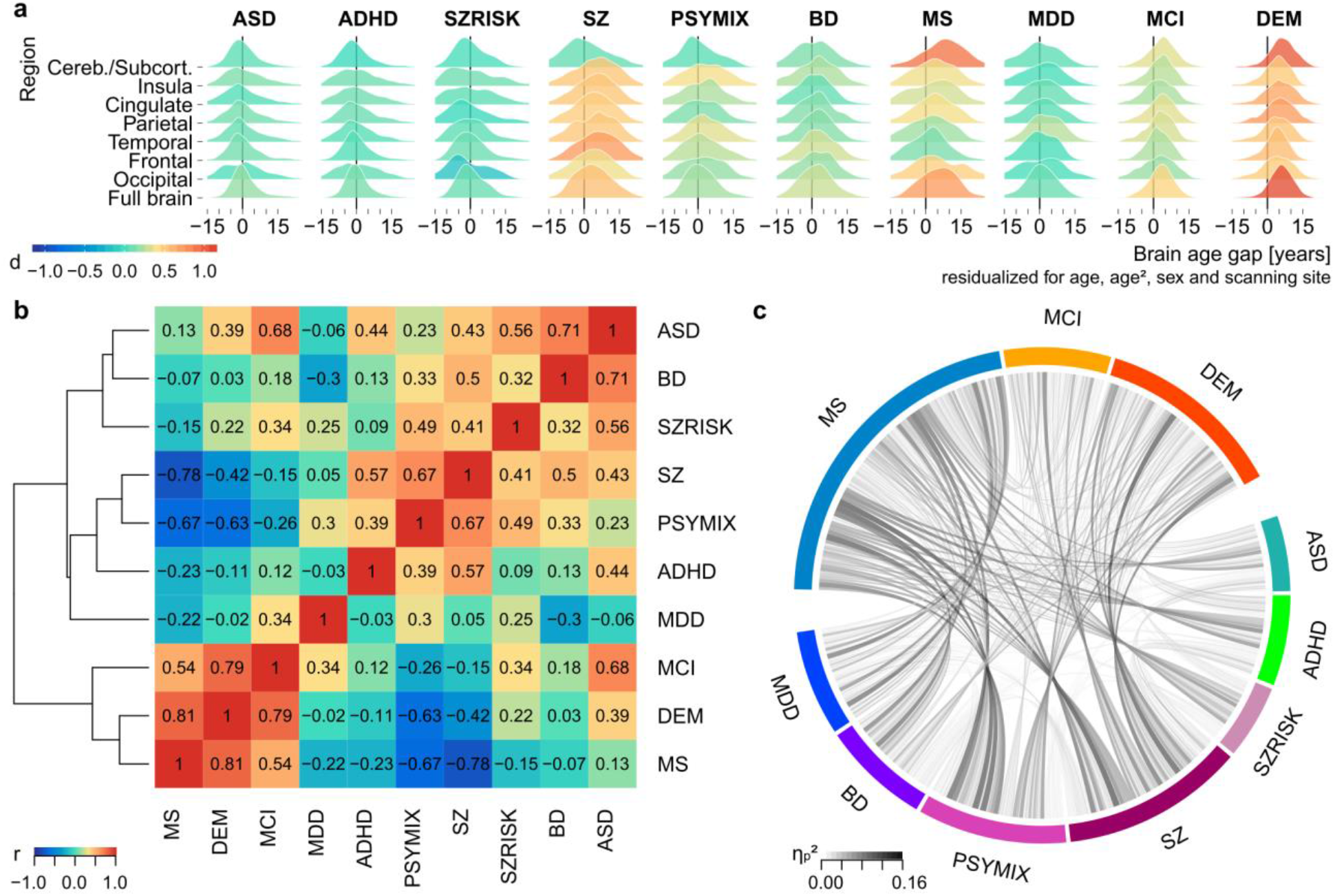
Several disorders displayed regional specific aging patterns. **a**, Region-wise brain age gaps per disorder. Colours indicate Cohen’s d effect sizes for group comparison to healthy controls matched for age, sex and scanning site. Strongest patterns were observed in the cerebellar/subcortical region for DEM and MS, and in the frontal lobe for SZ. **b**, Correlation matrix based on the effect sizes from panel (a) indicates similarities (e.g. between MS, DEM and MCI) and dissimilarities (e.g. between SZ/PSYMIX and MS/DEM) in the spatial brain age patterns between the disorders. Sorting is based on hierarchical clustering. **c**, Effect sizes of region x group interaction effects from repeated measures ANOVAs run for each combination of regions and groups (1260 tests in total). Strongest interaction effects were observed between MS/MCI/DEM and the other disorders, confirming the dissimilarity pattern from panel (b) with an independent analysis.

Figure 3b illustrates a hierarchical clustering of clinical groups based on region-wise effect sizes. One cluster of groups with similar spatial brain age gap patterns comprised MS, MCI and DEM, whereas the other groups formed a second cluster. Notably, the spatial patterns of the groups in the first cluster were negatively associated with several disorders in the second cluster, pointing toward spatial specificity of these disorders.

To explore these differences further, we tested for group x region interactions on each pairwise combination of clinical groups and pairwise combination of region-wise brain age gaps (1260 tests). Figure 3c illustrates the effect sizes for all resulting group x region interactions. Confirming the results from Figure 3b, strongest interaction effects were observed between the groups from cluster 1 and those from cluster 2. For example, the three strongest interaction effects indicated that brain age gaps for frontal and cerebellar/subcortical regions diverged mostly between MS and SZ (Partial Eta squared *η*_*p*_*²* = 0.16), MS and PSYMIX (*η*_*p*_*²* = 0.15) and between SZ and DEM (*η*_*p*_*²* =0.15; Suppl. Fig. 5 provides effect sizes for all tests). Together, these results demonstrate that several common disorders affecting the brain show anatomically differential patterns of increased brain age gap, indicating that the rate at which different regions age in relation to each other oftentimes showed opposite patterns between disorders typically considered neurodevelopmental and neurodegenerative disorders, respectively.

With converging findings suggesting largest brain age gaps in SZ, MS, MCI and DEM, we explored the functional relevance of the region-wise brain age gaps for these groups, testing for associations with clinical and cognitive data. Clinical data available in the SZ test sample included symptom (*n* = 81 HC, *n* = 391 SZ) and function (*n* = 271 SZ) scores of the Global Assessment of Functioning scale^25^ (GAF) as well as positive (*n* = 57 HC, *n* = 653 SZ) and negative (*n* = 57 HC, *n* = 655 SZ) scores of the Positive and Negative Syndrome Scale^26^ (PANSS). In the MS test sample, we assessed associations with scores from the Expanded Disability Status Scale^27^ (EDSS, *n* = 188 MS) and in the joint MCI and DEM test samples, we assessed associations with Mini Mental State Examination scores^28^ (MMSE, *n* = 901 HC, *n* = 921 MCI, *n* = 707 DEM). Figure 4a depicts association strengths for all tests and Figure 4b illustrates the strongest association for each test, except for the PANSS scores where only weak associations were found. In SZ, larger brain age gaps were associated with lower functioning, in particular for full brain, frontal, temporal and insula brain age gaps (GAF function all z < −0.18, all *P* < 0.003; GAF symptom all *z* < −0.20, all *P* < 2 × 10^−5^). In MS, larger brain age gap was associated with higher disability, in particular for the full brain age gap (*z* = 0.23, *P* = .001). Finally, lower cognitive functioning was associated with larger brain age gaps in the joint MCI/DEM samples, with strongest effects for full brain (*z* = −0.34, *P* = 4 × 10^−65^) and cerebellar/subcortical (z = −0.31, *P* = 2 × 10^−53^) brain age gaps.

**Figure 4.**
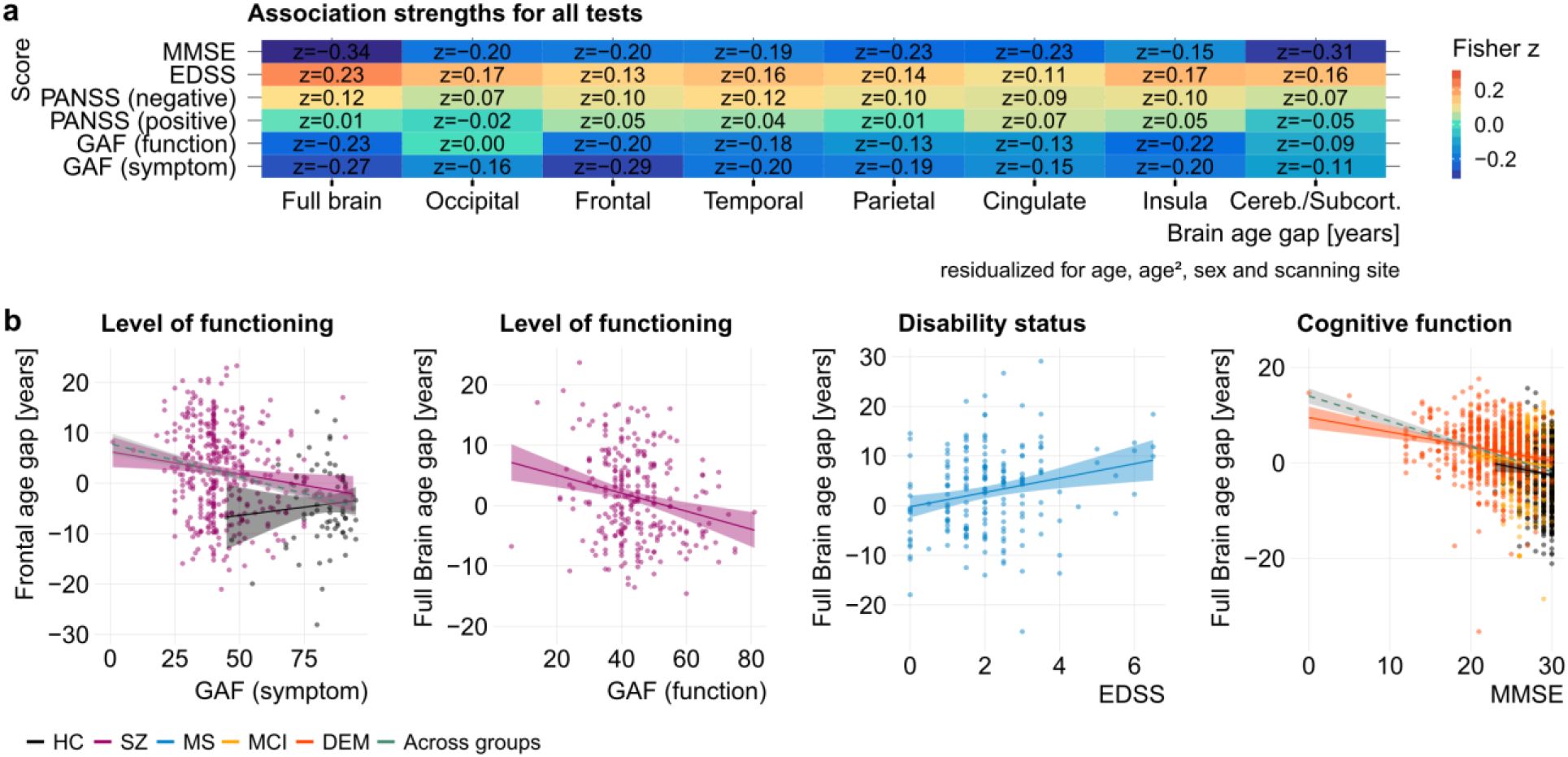
Region-wise brain age gaps were associated with cognitive and clinical scores. **a**, Fisher z score for all tested associations. In the SZ test sample, we assessed associations with GAF (function and symptom) and PANSS (positive and negative). In MS, we tested for associations with EDSS and in the joint MCI and DEM test samples, we assessed associations with MMSE scores. **b**, Exemplary illustration of the strongest associations for GAF, EDSS and MMSE.

## The genetic architecture of brain age gap

Given the known genetic contributions to brain disorders, our results pose the question to what degree brain age patterns are genetically constrained and if the implied genes overlap with the polygenic architectures of the disorders. In our cohorts, single nucleotide polymorphism (SNP) data were available for 16,269 adult healthy controls with European ancestry, after disregarding subsets with data from clinical groups, children and individuals with non-European ancestry, all of which were too small to warrant an analysis. We estimated full and region-wise brain age for these individuals using 5-fold cross-validation in a model trained on all healthy controls (*n* = 30,967) and regressed age, age², sex, and scanning site effects from the resulting brain age gaps.

First, we ran genome-wide complex trait analysis (GCTA^29^) across all 16,269 individuals, including the first four population components from multidimensional scaling as covariates to further control for population stratification. The results revealed significant heritability (Fig. 5a), with common SNPs explaining 18.3% of the variance in brain age gap across all individuals (full brain, h^2^_SNP_ = 0.1828, SE = 0.02, *P* < 1 × 10^−16^) and 11.1-18.4% of the variance in region-wise brain age gaps (all *P* < 2 × 10^−9^).

**Fig. 5:**
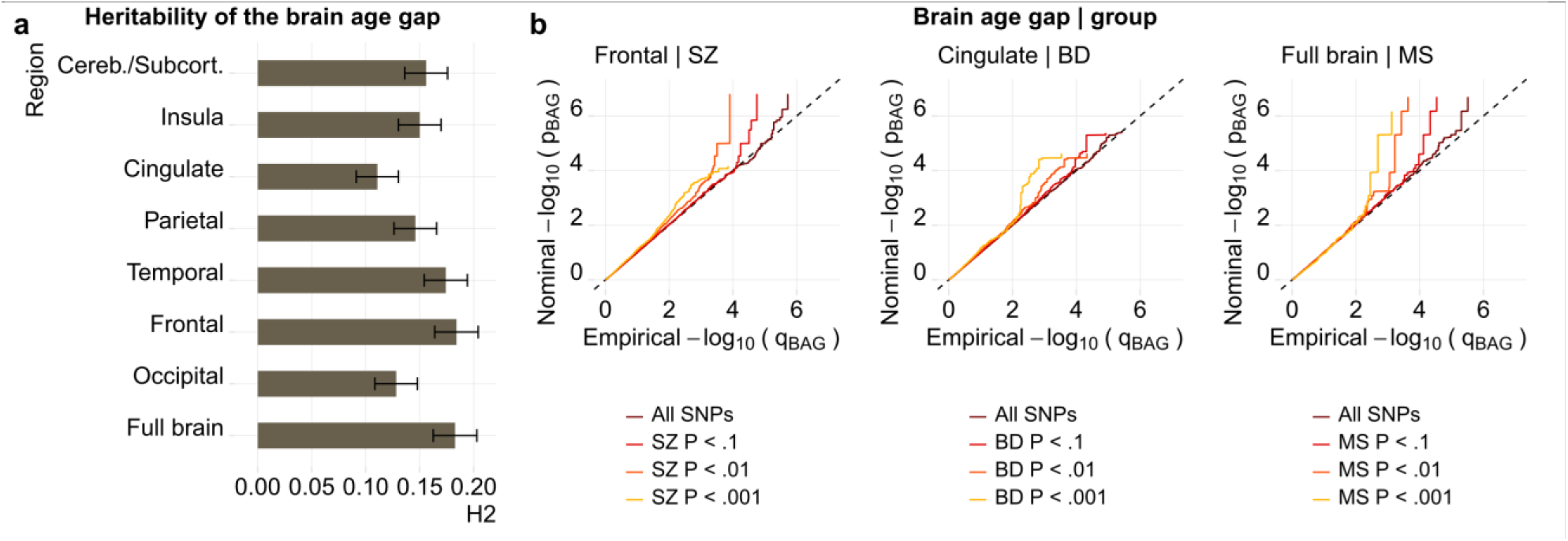
The brain age gaps are heritable and the genetic underpinnings overlap with those observed for several disorders. **a**, Heritability (H2) estimated using GCTA (all *P* < 2 × 10^−9^). **b**, Exemplary illustration of genetic enrichment between brain age gaps and SZ, BD and MS, assessed using conditional Q-Q plots. The dashed line is the expected trajectory under the global null hypothesis, whereas the coloured lines are the trajectories observed in the complete set, and in subsets of SNPs identified by their association with the disorder. Abbreviations: BAG, brain age gap. SNP, single nucleotide polymorphism.

Next, we assessed the overlap between the genetic underpinnings of brain age gap and common brain disorders. Focusing on those disorders that showed a significant brain age gap in the main analysis, we gathered genome-wide association analysis (GWAS) summary statistics for SZ and BD from the Psychiatric Genomics Consortium^30,31^, MS from the International Multiple Sclerosis Genetics Consortium^32^, and AD from the International Genomics of Alzheimer’s Project^33^. In addition, we performed GWAS on the full brain and region-wise brain age gaps in the above-described set of 16,269 healthy controls. We used conditional Q-Q plots^34^ to assess polygenic overlap between two complex traits, conditioning GWAS summary statistics from each of the brain age gaps on GWAS summary statistics from each of the disorders. Notably, our results indicate genetic overlap between brain age gap and brain disorders. Figure 5b provides exemplary illustrations of conditional Q-Q plots for the frontal brain age gap stratified by SZ, the cingulate brain age gap stratified by BD and the full brain age gap stratified by MS. When selecting subsets of SNPs based on their associations with the disorders, the nominal −log10 transformed P-values of the brain age gaps deviated from the trajectories expected under the global null hypothesis, indicating that the brain age gaps are enriched for SNP associations with the relevant disorder. SZ and MS also showed patterns of enrichment with subcortical brain age gap and BD with frontal brain age gap, whereas no clear patterns were observed for AD (Suppl. Fig. 6).

Next, we combined GWAS summary statistics of brain age gaps and the disorders in conjunctional FDR analyses^35,36^, to identify SNPs that are associated with both phenotypes. We found 15 independent, significant loci showing pleiotropy between brain age gaps and SZ (2 occipital, 4 frontal, 3 temporal, 1 parietal, 2 cingulate, 1 insula, 2 cerebellar/subcortical; 116 SNPs in total), 6 loci for BD (3 frontal, 2 cingulate, 1 insula; 40 SNPs in total), 7 loci for MS (2 full brain, 2 frontal, 1 temporal, 2 subcortical; 7 SNPs in total) and 1 locus for AD (temporal, 1 SNP), respectively (Suppl. Table 3). An intronic variant (rs940904) in protein coding gene PITPNM2 at chromosome 12q24.31 underlying the frontal brain age gap significantly overlapped both with SZ and MS.

## Discussion

Taken together, our results provide strong evidence that several common brain disorders are associated with accentuated aging of the brain compared to chronological age, with effects observed in SZ, PSYMIX, BD, MS, MDD, MCI and DEM; but not in ASD, ADHD or SZRISK. Importantly, we revealed a distinct neuroanatomical distribution of brain age gaps in several disorders. Associations with clinical and cognitive data underlined the functional relevance of the brain age gaps and genetic analyses in healthy controls provided evidence that the brain age gaps are heritable, with overlapping genes implicated in the genetic underpinnings of brain age gaps and common brain disorders.

Our approach of estimating brain age at the level of brain regions was useful to reveal differential spatial patterns between disorders. Whereas the implicated regions in the spatial brain age profiles of the disorders matched previously reported structural and functional abnormalities (e.g. frontal in SZ^37-39^, or the widespread volume loss in AD with large effects in subcortical structures^40^), our region-wise brain age approach preserved the well-established benefit of down-sampling a large number of brain imaging features into a highly condensed and interpretable score without a total loss of spatial sensitivity. As such, the analysis revealed substantial differences in spatial aging profiles between disorders typically regarded neurodegenerative disorders (MS, MCI, DEM) and disorders with established neurodevelopmental sources, especially SZ and PSYMIX. Whereas these disorders were all associated with an increased brain age gap on the full brain level, the region-wise analysis uncovered an interaction between the frontal brain age patterns observed in SZ and PSYMIX and the cerebellar/subcortical patterns observed in MS and DEM. Moreover, brain age gaps covered functional relevance beyond the group differences. We identified significant associations with clinical and cognitive data, in particular with scores of the Global Assessment of Functioning scale^25^ in SZ, with the Expanded Disability Status Scale^27^ in MS and with Mini Mental State Examination scores^28^ in the dementia spectrum. These results may warrant further research in which the link between the rate of changes in brain age gaps and the clinical and cognitive outcome can be studied in a longitudinal setting. Depending on the sensitivity of such associations to dynamic changes in clinical and cognitive function, such studies may also explore the biological mechanisms underlying these associations as a potential anchor point for treatment options.

Genetic analysis offers one way of exploring the constraining factors underlying phenotypic variation. Here, we provided evidence that full and region-wise brain age gaps represent genetically influenced traits, and illustrated that the genetic variants associated with brain age gaps are also associated with SZ, BD, MS and AD. In line with the accumulating evidence that common disorders of the brain are highly polygenic and partly overlapping^30-34,36,41^, these results suggest shared molecular genetic mechanisms between brain age gaps and brain disorders. Statistical associations do not necessarily signify causation, and functional interpretations of the identified genes should be made with caution. Larger imaging genetics samples, in particular those including individuals with common brain disorders, may in the future allow the investigation of specificity of the implicated genes. Considering the observed interaction effects in spatial brain age profiles between some disorders, we speculate that such analyses may offer novel insight into specific molecular mechanisms and will allow us to delineate the processes that affect the pace and profile of global and regional brain aging for each of these disorders.

In conclusion, in this largest brain age study to date, we established that the brain age gap is genetically constrained, increased in several common brain disorders, and linked to clinical and cognitive phenotypes. Our results establish the potential of advanced lifespan modelling in the clinical neurosciences, highlighting the benefit of big data resources that cover a wide span of ages and disease conditions. Delineating dynamic lifespan trajectories within and across individuals will be essential to disentangle the pathophysiological complexity of brain disorders.

## Acknowledgements

The authors were funded by the Research Council of Norway (213837, 223273, 204966/F20, 229129, 249795/F20, 276082), the South-Eastern Norway Regional Health Authority (2013-123, 2014-097, 2015-073), Stiftelsen Kristian Gerhard Jebsen, and the European Commission 7th Framework Programme (602450, IMAGEMEND). The data used in this study were gathered from various sources. A detailed overview of the included cohorts and acknowledgement of their respective funding sources and cohort-specific details is provided in Supplementary Table 1.

## Author contributions

T.K. and L.T.W. conceived the study; T.K., N.T.D. and L.T.W. pre-processed all data in Freesurfer; N.T.D., M.J.L., C.L.B, L.B.N., L.T.W. and T.K. performed quality control of the data; T.K. performed the analysis with contributions from L.T.W. and D.v.d.M.; T.K., L.T.W., N.T.D., D.v.d.M. and O.A.A. contributed to interpretation of the results. All remaining authors were involved in data collection at various sites as well as cohort-specific tasks. T.K. and L.T.W. wrote the first draft of the paper and all authors contributed to and approved the final manuscript.

## Materials & Correspondence

The data incorporated in this work were gathered from various resources (see acknowledgements). Material requests will need to be placed with individual PIs. Corresponding authors Tobias Kaufmann and Lars T. Westlye will provide additional detail upon correspondence.

## Collaborators

Members of the Karolinska Schizophrenia Project (KaSP): Farde L^1^, Flyckt L^1^, Engberg G^2^, Erhardt S^2^, Fatouros-Bergman H^1^, Cervenka S^1^, Schwieler L^2^, Piehl F^3^, Agartz I^1,4,5^, Collste K^1^, Victorsson P^1^, Malmqvist A^2^, Hedberg M^2^, Orhan F^2^

^1^Centre for Psychiatry Research, Department of Clinical Neuroscience, Karolinska Institutet, & Stockholm County Council, Stockholm, Sweden; ^2^Department of Physiology and Pharmacology, Karolinska Institutet, Stockholm, Sweden; ^3^Neuroimmunology Unit, Department of Clinical Neuroscience, Karolinska Institutet, Stockholm, Sweden; ^4^NORMENT, KG Jebsen Centre for Psychosis Research, Division of Mental Health and Addiction, University of Oslo, Oslo, Norway; ^5^Department of Psychiatry Research, Diakonhjemmet Hospital, Oslo, Norway.

## Online methods

### Samples

We have included data collected through collaborations, data sharing platforms, consortia as well as available in-house cohorts. Supplementary Table 1 and 2 provide detailed information on the individual cohorts. All included cohorts have been published on, and we refer to a list of publications that can be consulted for a more detailed overview of cohort characteristics. Data collection in each cohort was performed with participants’ written informed consent and with approval by the respective local Institutional Review Boards.

### Image pre-processing and quality control

Raw T1 data for all study participants were stored and analysed locally at University of Oslo, following a harmonized analysis protocol applied to each individual subject data (Suppl. Fig. 1). We performed automated surface-based morphometry and subcortical segmentation using Freesurfer 5.3^19^. We deployed an automated quality control protocol executed within each of the contributing cohorts (Suppl. Tables 1–2), that excluded potential outliers based on global data quality measures. In brief, we regressed age, age², sex and (in case of multiple scanners) scanning site from mean cortical thickness, cortex volume, subcortical grey matter volume and from estimated total intracranial volume. Next, we z-standardized the resulting absolute of the residuals and excluded those subjects that exceeded a pre-defined standard deviation (SD) threshold of 4 SD. On top of this, a random set of data was carefully screened by trained research personnel to identify segmentation errors, assess the quality of each subject’s brain images manually, edit segmentation where possible and to exclude data of insufficient quality (*n* = 3957 manually controlled, *n* = 166 excluded, *n* = 219 edited). Taken together, the main analysis excluded cases identified by manual QC as well as cases exceeding a threshold of 4 SD in the automated quality control on either of the four brain imaging measures, yielding 36,891 subjects. In addition, we performed supplementary analyses using a subset of data, where a more stringent quality control and exclusion procedure was applied. Using multivariate outlier detection based on robust methods as implemented in the R package mvoutlier^24^, we identified an additional 5513 data sets with potentially less sufficient data quality. Thus, supplemental analysis provides a sanity check with those subjects excluded (sample size: *n* = 31,378).

### Brain age prediction

We utilized the most recent cortical parcellation scheme^21^ to extract cortical thickness, area and volume for 180 regions of interest (ROI) per hemisphere. In addition, we extracted the classic set of cerebellar/subcortical and cortical summary statistics^19^. This yielded a total set of 1118 structural brain imaging features (360/360/360/38 for cortical thickness/area/volume as well as cerebellar/subcortical and cortical summary statistics, respectively).

We used machine learning on this feature set to predict the age of each individual’s brain. First, we split the available data into a training sample and ten independent test samples, as described in the main text (Fig. 1a). Next, for each sex, we trained machine learning models utilizing the *xgboost* package in R^42^, chosen due to its resource efficiency and demonstrated superior performance in previous machine learning competitions, to predict the age of the brain using data available in the training set. First, model parameters were tuned using a 5-fold cross-validation of the training data. This step identified the optimal number of model training iterations by assessing the prediction error for 1500 rounds and implementing an early stopping if the performance did not improve for 20 rounds. Based on previous experience, the learning rate was pre-set to *eta*=0.01 and all other parameters were set to default^42^ for linear *xgboost* tree models. After determining the optimal number of training iterations, the full set of training data was used to train the final models with the adjusted *nrounds* parameter. In addition, to assess overall model performance, prediction models were cross-validated within the training set using a 5-fold cross validation, each fold implementing the above described training procedure and testing on the hold-out part of the training set. Brain age predictions on the level of individual brain regions followed the same procedures as those described for the full brain level, except that the feature set was reduced to cover only those features that overlapped more than 50% with a given lobe. Regions were defined following the Freesurfer *lobesStrict* segmentation as *occipital*, *frontal*, *temporal*, *parietal*, *cingulate* and *insula*. In addition, given the limited number of cerebellar features available in the Freesurfer summary statistics, cerebellar and subcortical features were grouped into a *cerebellar*/*subcortical* region (Fig. 1b).

For the genetic analyses, a different approach had to be taken to ensure maximum exploitation of the limited availability of genetic data. Rather than splitting into training and test sets, we selected all healthy subjects and estimated their brain age using a 5-fold cross-validation approach like the one performed when validating performance of the training set. The resulting unbiased estimates of brain age gaps for all individuals with genetic data available went into the genome-wide complex trait analysis and conjunctional FDR.

### Main statistical analysis framework

We performed both mega- (across cohorts) and meta- (within cohort) analyses. To estimate group effects on a given measure in a mega-analysis framework, we computed the effect of diagnosis in relation to the healthy controls for each of the ten test samples in a linear model accounting for age, age², sex and scanning site. Cohen’s d effect sizes were estimated based on contrast t-statistics^43^ following Formula 1:

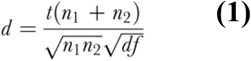

For the meta-analysis, similar models were computed within cohorts. In addition to estimating Cohen’s d (Formula 1), we estimated the variance of d following Formula 2.

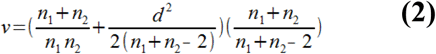

Cumulative effects across cohorts were then estimated using a variance-weighted random-effects model as implemented in the *metafor* package in R^44^.

### Assessment of regional specificity

The clustering in Figure 3b was performed using heatmap.2 from the *gplots* package^45^ in R. A correlation matrix was computed based on the case-control effect sizes obtained from each test sample and region and hierarchical clustering was performed using the default settings. To further explore regional specificity, we performed another analysis that involved only the clinical groups. We regressed age, age², sex and scanning site from the brain age gaps in each test sample. Next, we joined data from each pair of clinical groups and each pair of regions for repeated measures analysis of variance and estimated the effect sizes of region x group interactions (1260 ANOVAs in total). The interaction effects were visualized in Figure 3c using the *circlize* package^46^ in R.

### Genetic analyses

We restricted all genetic analyses to individuals with European ancestry, as determined through multidimensional scaling (MDS), and included the first four population components as covariates to further control for population stratification. Single Nucleotide Polymorphism (SNP) data were available for 16,269 adult healthy individuals with European ancestry. We used genome-wide complex trait analysis^29^ (GCTA) to estimate the proportion of variance in brain age explained by SNPs. Before the analysis, we removed high LD regions from the genetic data and pruned it, using a sliding window approach with a window size of 50 base pair (bp), a step size of 5 bp and an r^2^ of 0.2, leaving 133,147 SNPs. All GCTA analyses accounted for age, age², sex, scanning site and genetic batch.

Furthermore, we used conditional Q-Q plots^34^ and conjunctional FDR analyses^35,36^ to assess polygenic overlap between two complex traits. We gathered genome-wide association analysis (GWAS) summary statistics for SZ and BD from the Psychiatric Genomics Consortium^30,31^, MS from the International Multiple Sclerosis Genetics Consortium^32^, and AD from the International Genomics of Alzheimer’s Project^33^; and performed GWAS on the full brain and region-wise brain age gaps in the above-described sample of 16,269 healthy adults. The MHC region was excluded from the analysis. The SNPs were pruned using a pairwise correlation coefficient approximation to LD (r²), where SNPs were disregarded at r²<0.2 and pruning performed with 20 iterations, as described elsewhere^34^. Conjunctional FDR was run for each pair of full brain / region-wise brain age gap and group, using conjunctional FDR threshold of 0.05. SNPs were annotated using the Ensembl Variant Effect Predictor^47^.

### Cognitive and clinical associations

Cognitive and clinical associations were tested in subsets based on data availability as described in the main text. First, we regressed age, age², sex and scanning site from the brain age gaps. Next, we correlated the resulting residuals with scores of the Global Assessment of Functioning scale^25^ (GAF), the Positive and Negative Syndrome Scale^26^ (PANSS), the Expanded Disability Status Scale^27^ (EDSS) and Mini Mental State Examination scores^28^ (MMSE). We transformed the resulting correlations using Fisher’s z transform. Therefore, the reported associations essentially reflect a partial correlation between full brain / region-wise brain age gaps and clinical/cognitive scores, controlling for confounding effects of age, sex and site.

## Code availability

The main analysis was performed using R statistics^48^. The code needed to reproduce the results is available from the authors upon request.

---

**Suppl. Figure 1:**
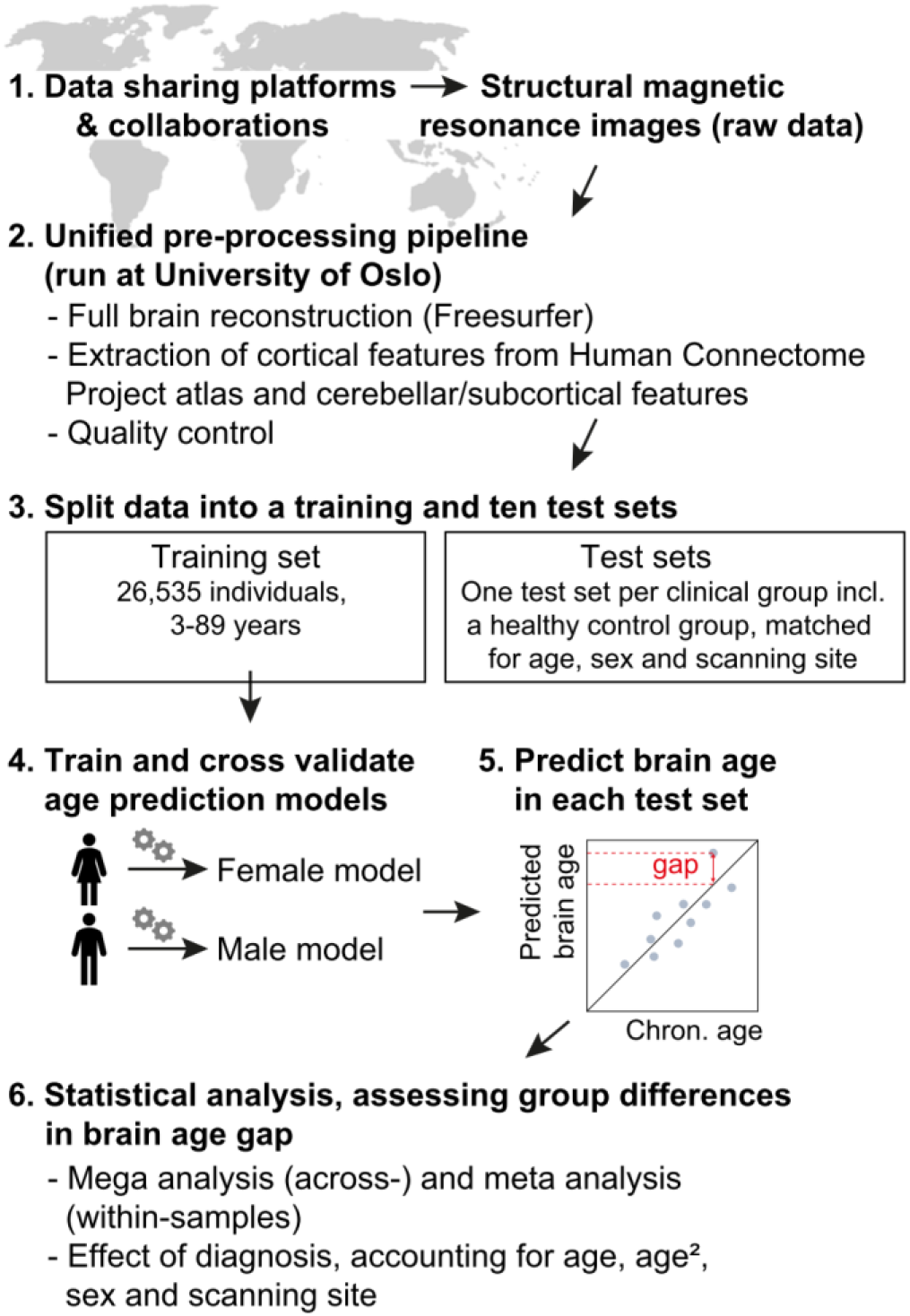
Outline of the main analysis pipeline.

**Suppl. Figure 2:**
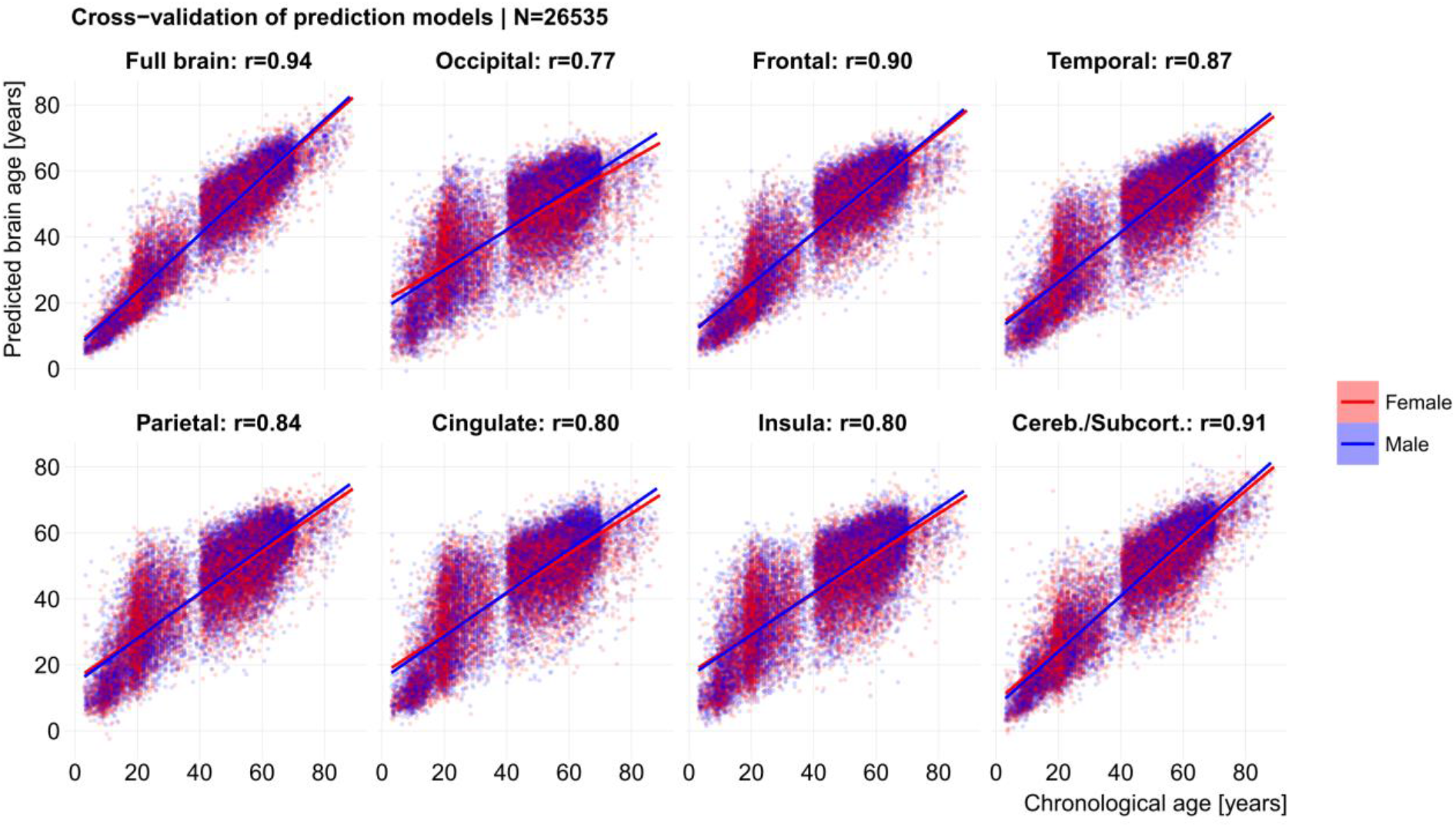
Cross-validation of the prediction models confirmed validity of the models. The correlation between chronological age and predicted brain age estimated using 5-fold cross-validation within the training set is shown for each feature set.

**Suppl. Figure 3:**
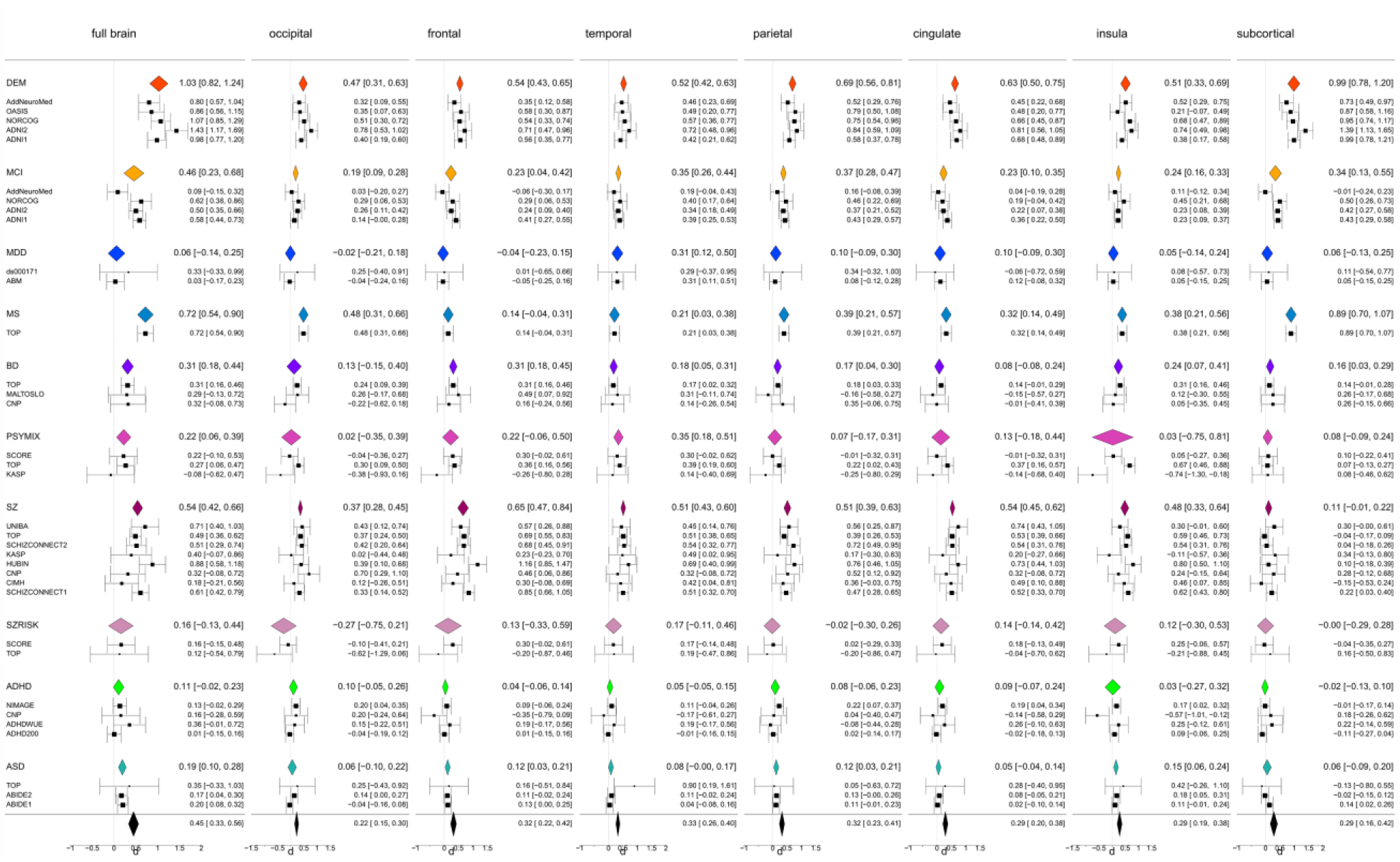
Meta-analysis confirmed mega-analysis results. All Cohen’s d effect sizes for the effect of group accounted for age, age² and sex. Further, Cohen’s d for all cohorts that were collected at multiple sites also accounted for scanning site.

**Suppl. Figure 4:**
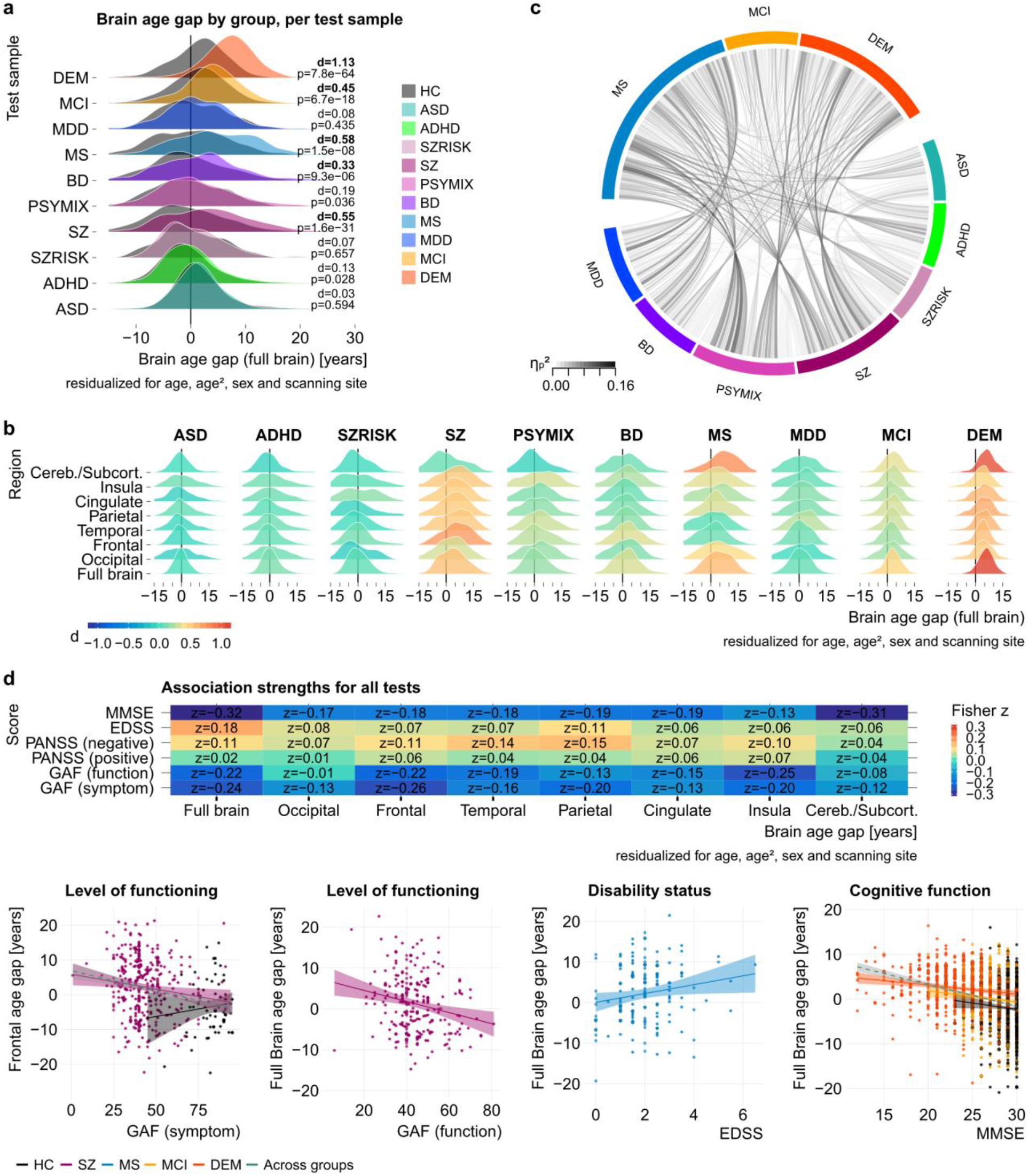
Replication of results in a subset of 31,378 individuals following more stringent, multivariate exclusion criteria. **a**, Replication of group effects (for comparison, see Fig. 2) **b**, Replication of spatial brain age gap patterns (for comparison, see Fig. 3a) **c**, Replication of interaction effect pattern (for comparison, see Fig 3c). **d**, Replication of associations with clinical and cognitive scores (for comparison, see Fig. 4).

**Suppl. Figure 5:**
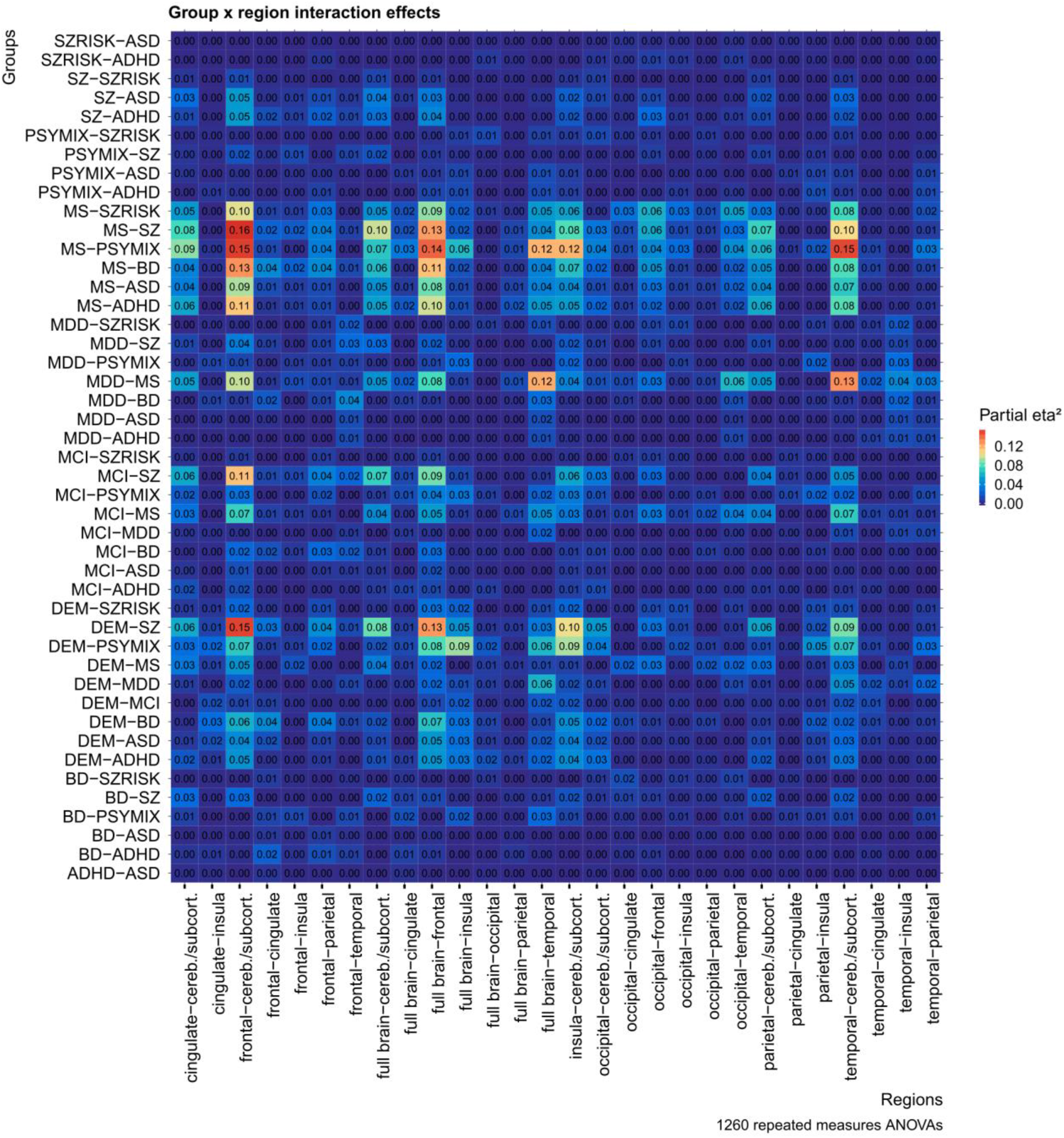
Results from 1260 repeated measures ANOVAs confirm group x region interaction effects in brain age patterns. Strongest effects were observed between MS and SZ, MS and PSYMIX as well as DEM and SZ, suggesting divergent aging patterns in these disorders.

**Suppl. Figure 6:**
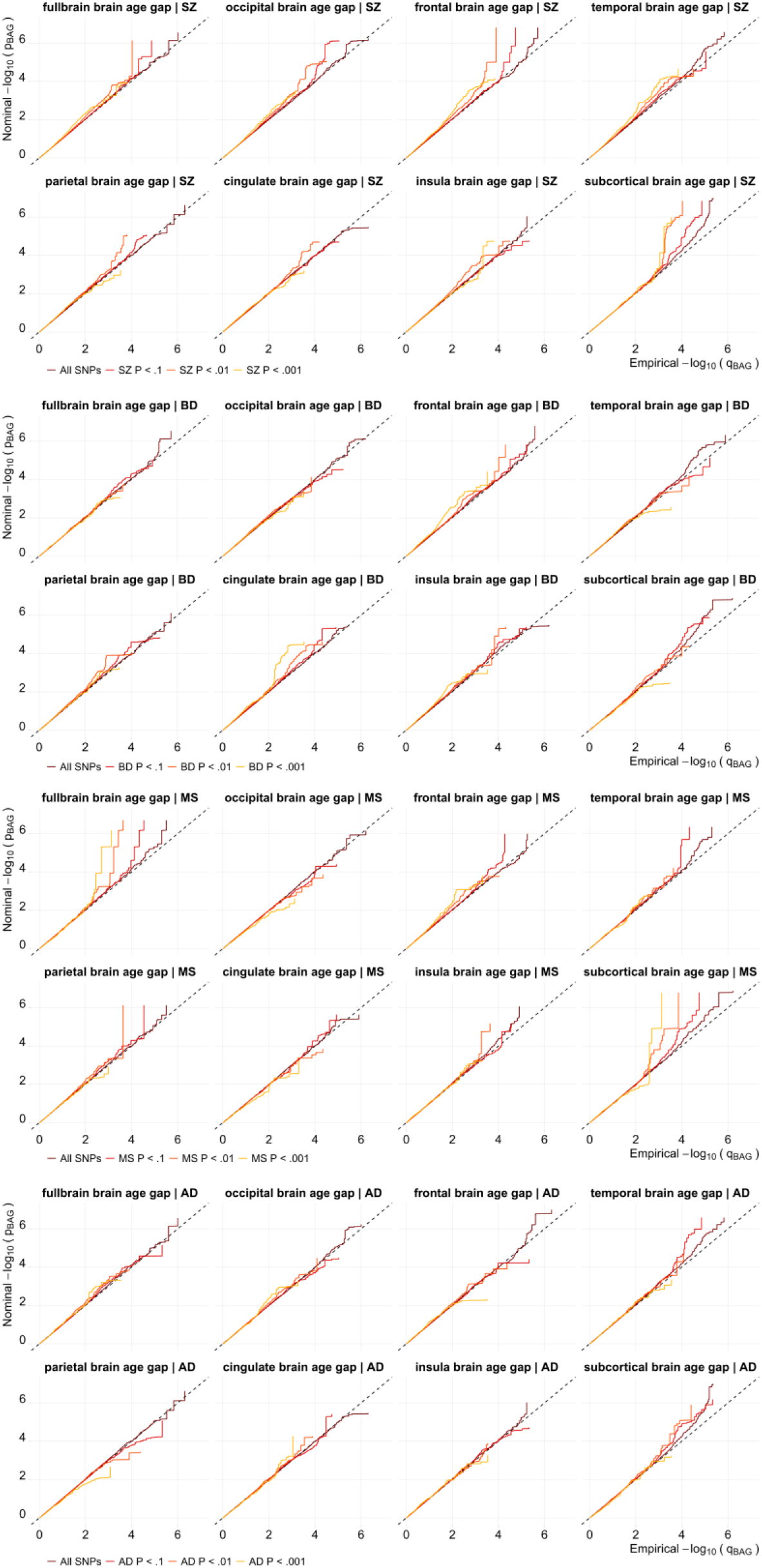
Conditional Q-Q plots for all brain age gaps conditioned for SZ, BD, MS or AD suggests enrichment for several brain age gaps.

**Suppl. Table 1:**
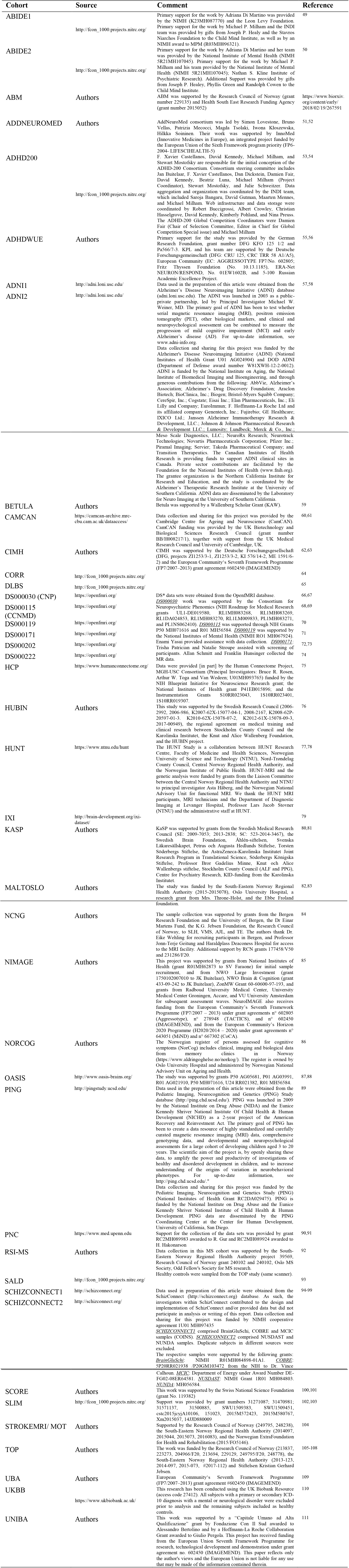
Summary of the included cohorts. Several of the cohorts are under ongoing data collection, thus the subject numbers provided in the reference publications may not match those in this article.

**Suppl. Table 2:**
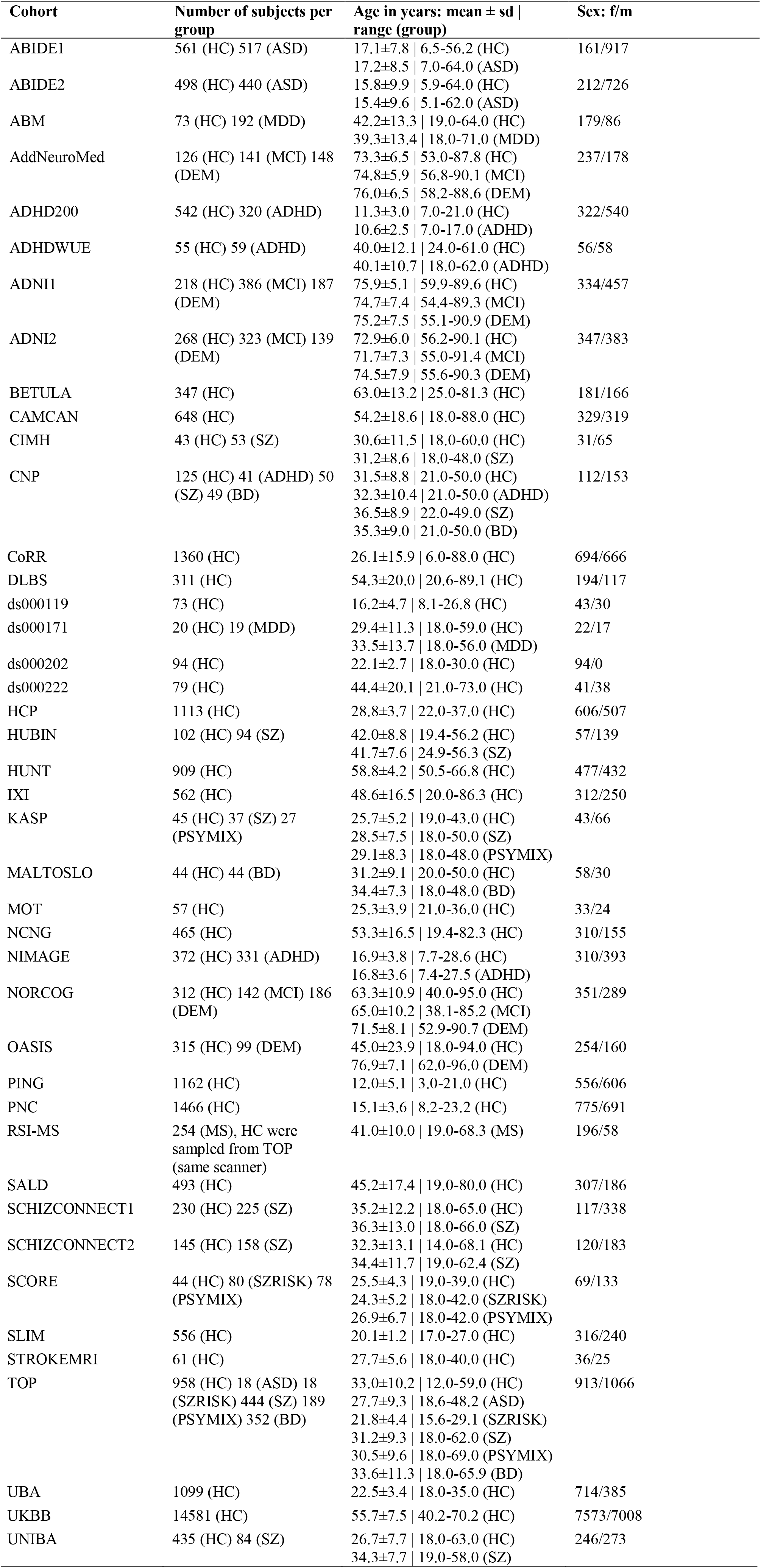
Summary of group size, age and sex for each cohort.

| Cohort | Number of subjects per group | Age in years: mean ± sd \| range (group) | Sex: f/m |
| --- | --- | --- | --- |
| ABIDE1 | 561 (HC) 517 (ASD) | 17.1±7.8 \| 6.5-56.2 (HC)<br>17.2±8.5 \| 7.0-64.0 (ASD) | 161/917 |
| ABIDE2 | 498 (HC) 440 (ASD) | 15.8±9.9 \| 5.9-64.0 (HC)<br>15.4±9.6 \| 5.1-62.0 (ASD) | 212/726 |
| ABM | 73 (HC) 192 (MDD) | 42.2±13.3 \| 19.0-64.0 (HC)<br>39.3±13.4 \| 18.0-71.0 (MDD) | 179/86 |
| AddNeuroMed | 126 (HC) 141 (MCI) 148 (DEM) | 73.3±6.5 \| 53.0-87.8 (HC)<br>74.8±5.9 \| 56.8-90.1 (MCI)<br>76.0±6.5 \| 58.2-88.6 (DEM) | 237/178 |
| ADHD200 | 542 (HC) 320 (ADHD) | 11.3±3.0 \| 7.0-21.0 (HC)<br>10.6±2.5 \| 7.0-17.0 (ADHD) | 322/540 |
| ADHDWUE | 55 (HC) 59 (ADHD) | 40.0±12.1 \| 24.0-61.0 (HC)<br>40.1±10.7 \| 18.0-62.0 (ADHD) | 56/58 |
| ADNI1 | 218 (HC) 386 (MCI) 187 (DEM) | 75.9±5.1 \| 59.9-89.6 (HC)<br>74.7±7.4 \| 54.4-89.3 (MCI)<br>75.2±7.5 \| 55.1-90.9 (DEM) | 334/457 |
| ADNI2 | 268 (HC) 323 (MCI) 139 (DEM) | 72.9±6.0 \| 56.2-90.1 (HC)<br>71.7±7.3 \| 55.0-91.4 (MCI)<br>74.5±7.9 \| 55.6-90.3 (DEM) | 347/383 |
| BETULA | 347 (HC) | 63.0±13.2 \| 25.0-81.3 (HC) | 181/166 |
| CAMCAN | 648 (HC) | 54.2±18.6 \| 18.0-88.0 (HC) | 329/319 |
| CIMH | 43 (HC) 53 (SZ) | 30.6±11.5 \| 18.0-60.0 (HC)<br>31.2±8.6 \| 18.0-48.0 (SZ) | 31/65 |
| CNP | 125 (HC) 41 (ADHD) 50 (SZ) 49 (BD) | 31.5±8.8 \| 21.0-50.0 (HC)<br>32.3±10.4 \| 21.0-50.0 (ADHD)<br>36.5±8.9 \| 22.0-49.0 (SZ)<br>35.3±9.0 \| 21.0-50.0 (BD) | 112/153 |
| CoRR | 1360 (HC) | 26.1±15.9 \| 6.0-88.0 (HC) | 694/666 |
| DLBS | 311 (HC) | 54.3±20.0 \| 20.6-89.1 (HC) | 194/117 |
| ds000119 | 73 (HC) | 16.2±4.7 \| 8.1-26.8 (HC) | 43/30 |
| ds000171 | 20 (HC) 19 (MDD) | 29.4±11.3 \| 18.0-59.0 (HC)<br>33.5±13.7 \| 18.0-56.0 (MDD) | 22/17 |
| ds000202 | 94 (HC) | 22.1±2.7 \| 18.0-30.0 (HC) | 94/0 |
| ds000222 | 79 (HC) | 44.4±20.1 \| 21.0-73.0 (HC) | 41/38 |
| HCP | 1113 (HC) | 28.8±3.7 \| 22.0-37.0 (HC) | 606/507 |
| HUBIN | 102 (HC) 94 (SZ) | 42.0±8.8 \| 19.4-56.2 (HC)<br>41.7±7.6 \| 24.9-56.3 (SZ) | 57/139 |
| HUNT | 909 (HC) | 58.8±4.2 \| 50.5-66.8 (HC) | 477/432 |
| IXI | 562 (HC) | 48.6±16.5 \| 20.0-86.3 (HC) | 312/250 |
| KASP | 45 (HC) 37 (SZ) 27 (PSYMIX) | 25.7±5.2 \| 19.0-43.0 (HC)<br>28.5±7.5 \| 18.0-50.0 (SZ)<br>29.1±8.3 \| 18.0-48.0 (PSYMIX) | 43/66 |
| MALTOSLO | 44 (HC) 44 (BD) | 31.2±9.1 \| 20.0-50.0 (HC)<br>34.4±7.3 \| 18.0-48.0 (BD) | 58/30 |
| MOT | 57 (HC) | 25.3±3.9 \| 21.0-36.0 (HC) | 33/24 |
| NCNG | 465 (HC) | 53.3±16.5 \| 19.4-82.3 (HC) | 310/155 |
| NIMAGE | 372 (HC) 331 (ADHD) | 16.9±3.8 \| 7.7-28.6 (HC)<br>16.8±3.6 \| 7.4-27.5 (ADHD) | 310/393 |
| NORCOG | 312 (HC) 142 (MCI) 186 (DEM) | 63.3±10.9 \| 40.0-95.0 (HC)<br>65.0±10.2 \| 38.1-85.2 (MCI)<br>71.5±8.1 \| 52.9-90.7 (DEM) | 351/289 |
| OASIS | 315 (HC) 99 (DEM) | 45.0±23.9 \| 18.0-94.0 (HC)<br>76.9±7.1 \| 62.0-96.0 (DEM) | 254/160 |
| PING | 1162 (HC) | 12.0±5.1 \| 3.0-21.0 (HC) | 556/606 |
| PNC | 1466 (HC) | 15.1±3.6 \| 8.2-23.2 (HC) | 775/691 |
| RSI-MS | 254 (MS), HC were sampled from TOP (same scanner) | 41.0±10.0 \| 19.0-68.3 (MS) | 196/58 |
| SALD | 493 (HC) | 45.2±17.4 \| 19.0-80.0 (HC) | 307/186 |
| SCHIZCONNECT1 | 230 (HC) 225 (SZ) | 35.2±12.2 \| 18.0-65.0 (HC)<br>36.3±13.0 \| 18.0-66.0 (SZ) | 117/338 |
| SCHIZCONNECT2 | 145 (HC) 158 (SZ) | 32.3±13.1 \| 14.0-68.1 (HC)<br>34.4±11.7 \| 19.0-62.4 (SZ) | 120/183 |
| SCORE | 44 (HC) 80 (SZRISK) 78 (PSYMIX) | 25.5±4.3 \| 19.0-39.0 (HC)<br>24.3±5.2 \| 18.0-42.0 (SZRISK)<br>26.9±6.7 \| 18.0-42.0 (PSYMIX) | 69/133 |
| SLIM | 556 (HC) | 20.1±1.2 \| 17.0-27.0 (HC) | 316/240 |
| STROKEMRI | 61 (HC) | 27.7±5.6 \| 18.0-40.0 (HC) | 36/25 |
| TOP | 958 (HC) 18 (ASD) 18 (SZRISK) 444 (SZ) 189 (PSYMIX) 352 (BD) | 33.0±10.2 \| 12.0-59.0 (HC)<br>27.7±9.3 \| 18.6-48.2 (ASD)<br>21.8±4.4 \| 15.6-29.1 (SZRISK)<br>31.2±9.3 \| 18.0-62.0 (SZ)<br>30.5±9.6 \| 18.0-69.0 (PSYMIX)<br>33.6±11.3 \| 18.0-65.9 (BD) | 913/1066 |
| UBA | 1099 (HC) | 22.5±3.4 \| 18.0-35.0 (HC) | 714/385 |
| UKBB | 14581 (HC) | 55.7±7.5 \| 40.2-70.2 (HC) | 7573/7008 |
| UNIBA | 435 (HC) 84 (SZ) | 26.7±7.7 \| 18.0-63.0 (HC)<br>34.3±7.7 \| 19.0-58.0 (SZ) | 246/273 |

**Suppl. Table 3:**
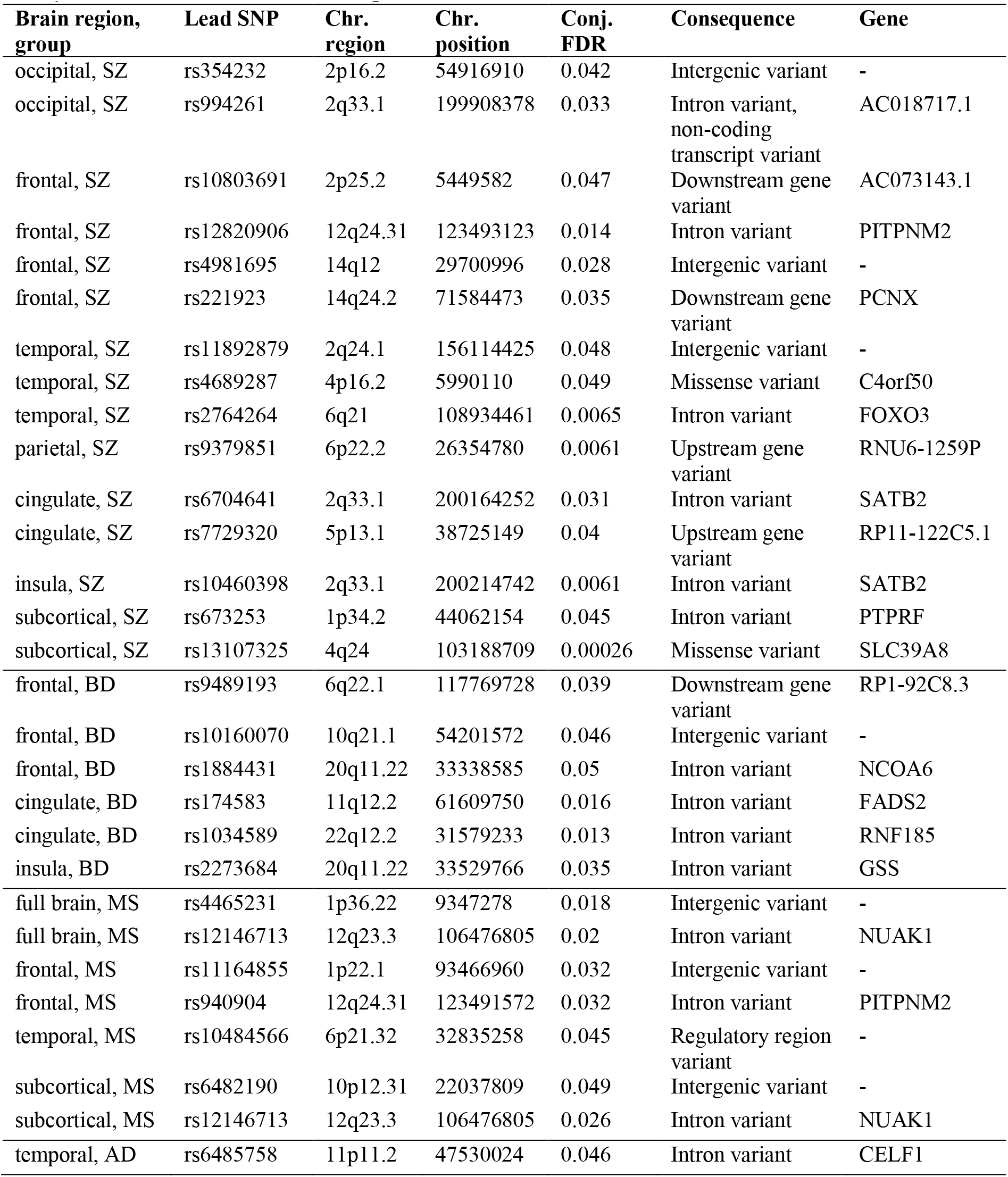
Significant loci from conjunctional FDR analysis reflecting overlap between brain age gaps and the respective disorders. The gene column reflects the gene closest to the significant SNP as identified via the Ensembl Variable Effect Predictor^47^, unless the closest gene is more than 5000 bp away in which case no annotation is provided.

